# Perilysosomal Ca2+ overload impairs autophagic degradation in β-cell lipotoxicity

**DOI:** 10.1101/2025.01.27.635047

**Authors:** Ha Thu Nguyen, Luong Dai Ly, Thuy Thi Thanh Ngo, Soo Kyung Lee, Carlos Noriega Polo, Subo Lee, Taesic Lee, Seung-Kuy Cha, Myung-Shik Lee, Andreas Wiederkehr, Kyu-Sang Park

**Author notes:** Correspondence: Claes B. Wollheim, M.D., Department of Cell Physiology and Metabolism, University Medical Center 1, rue Michel-Servet, CH-1211 Geneva 4, Switzerland, Kyu-Sang Park, M.D., Ph.D., Department of Physiology, Mitohormesis Research Center, Yonsei University Wonju College of Medicine, Wonju, 26426, Republic of Korea.

## Abstract

Saturated fatty acids impose lipotoxic stress on pancreatic β-cells, leading to β-cell failure and diabetes. In this study, we investigate the critical role of organellar Ca^2+^ disturbance on defective autophagy and β-cell lipotoxicity. Palmitate, a saturated fatty acid, induced perilysosomal Ca^2+^ elevation, sustained mTORC1 activation on the lysosomal membrane, suppression of the lysosomal transient receptor potential mucolipin 1 (TRPML1) channel, and accumulation of undigested autophagosomes in β-cells. These Ca^2+^ aberrations with autophagy defects by palmitate were prevented by a mTORC1 inhibitor or a mitochondrial superoxide scavenger. To alleviate perilysosomal Ca^2+^ overload, strategies such as lowering extracellular Ca^2+^, employing voltage-gated Ca^2+^ channel blocker or ATP-sensitive K^+^ channel opener effectively abrogated mTORC1 activation and preserved autophagy. Furthermore, redirecting perilysosomal Ca^2+^ into the endoplasmic reticulum (ER) with an ER Ca^2+^ ATPase activator, restores TRPML1 activity, promotes autophagic flux, and improves survival of β-cells exposed to palmitate-induced lipotoxicity. Our findings suggest oxidative stress-Ca^2+^ overload-mTORC1 pathway involves in TRPML1 suppression and defective autophagy during β-cell lipotoxicity. Restoring perilysosomal Ca^2+^ homeostasis emerges as a promising therapeutic strategy for metabolic diseases.

## INTRODUCTION

Type 2 diabetes (T2D) presents as a multifaceted metabolic disorder characterized by hyperglycemia resulting from impaired insulin secretion and insulin resistance(DeFronzo *et al*, 2015). Overnutrition and physical inactivity precipitate insulin resistance requiring increased insulin secretion to maintain glucose homeostasis. The capacity to adapt β-cell mass and insulin secretion is genetically determined, and β-cell decompensation leads to T2D in susceptible subjects(Alejandro *et al*, 2015; Nguyen *et al*, 2024). Within the diabetic milieu, free fatty acids are recognized as contributors to β-cell failure through a process known as lipotoxicity(Schaffer, 2003). Excess of saturated fatty acids initiate a cellular cascade, including oxidative stress, endoplasmic reticulum (ER) stress, inflammation, ultimately compromising the functionality and survival of β-cells(Nguyen *et al*., 2024).

Growing evidence suggests that impaired autophagic flux is linked to cell death induced by lipotoxicity(Las *et al*, 2011; Masini *et al*, 2009). Autophagy is an evolutionary conserved cellular process responsible for the degradation and recycling of damaged organelles, serving as a vital quality control mechanism to promote cell survival under various stress conditions. Upon lipotoxic stress, autophagy could act as a protective mechanism for β-cell survival by eliminating accumulated lipids and damaged organelles. However, chronic exposure to lipid excess, such as in obesity or elevated free fatty acid levels, can lead to an impaired and dysfunctional autophagic machinery(Choi *et al*, 2009; Cnop *et al*, 2014; Masini *et al*., 2009). One potential mechanism by which lipotoxicity impairs autophagy in β-cells is through upregulation of the mechanistic target of rapamycin complex 1 (mTORC1) signaling, which negatively regulates autophagic flux. Consistently, suppression of mTORC1 recovered insulin secretion in islets from T2D subjects which had higher expression level of mTORC1 than those from the controls(Yuan *et al*, 2017). However, the precise role of chronic mTORC1 activation by lipotoxic stress remains unclear.

Autophagy involves multiple processes, including the initiation, elongation, and maturation of autophagosomes, fusion between autophagosomes and lysosomes, and subsequent degradation of the materials(Meszaros *et al*, 2018). Lipotoxicity has been shown to disrupt the autophagosome-lysosome fusion, which is essential to complete the autophagic process(Guo *et al*, 2021; Liu *et al*, 2022; Mushtaq *et al*, 2023). This disruption hinders proper degradation, leading to the accumulation of toxic intermediates and damaged organelles. Previous studies suggest that the release of Ca^2+^ from lysosomes can dephosphorylate the transcription factor EB (TFEB), promoting its translocation to the nucleus, where it stimulates the induction of autophagy and lysosome biogenesis. Furthermore, lysosomal Ca^2+^ release triggers the biogenesis of autophagic vesicles through the generation of phosphatidylinositol 3-phosphate (PI3P) and the recruitment of essential PI3P-binding proteins to the nascent phagophore in a TFEB-independent manner(Scotto Rosato *et al*, 2019). Among the various types of Ca^2+^ channels in lysosomes, the mammalian mucolipin subfamily of transient receptor potential (TRPML) channels, also known as mucolipin TRP cation channel 1 (MCOLN), has been identified as particularly critical for autophagic flux(Zeevi *et al*, 2010). Additionally, Ca^2+^ release from the ER or lysosome is required for autophagosome-lysosome fusion, evidenced by the defective vesicular fusion with non-functional mutants of MCOLN1(Vergarajauregui *et al*, 2008) or through thapsigargin treatment(Ganley *et al*, 2011). It has been shown that mitochondrial reactive oxygen species (ROS) directly stimulate TRPML1 to release Ca^2+^. This triggers degradation of dysfunctional ROS-generating mitochondria, a quality control mechanism termed mitophagy(Zhang *et al*, 2016).

Several studies have clarified the role of cellular Ca^2+^ homeostasis in pancreatic β-cell physiology and its dysregulation in lipotoxicity(Ly *et al*, 2017). Disturbances in the regulation of ER, mitochondrial, and cytosolic Ca^2+^ due to an excess of saturated fatty acids lead to insulin secretory defects and cytotoxicity(Ly *et al*., 2017). In this context, we introduce a novel concept proposing that peri-lysosomal Ca^2+^ overload in β-cells, caused by sustained palmitate exposure, disrupts the proper function of TRPML1-mediated Ca^2+^ transients required for autophago-lysosomal degradation. We investigated molecular mechanisms of organellar Ca^2+^ dysregulation and its detrimental consequences for autophagic flux in palmitate-exposed β-cells. In addition, we explored novel therapeutic strategy aimed to recover perilysosomal Ca^2+^ homeostasis to alleviate lipotoxic stress and preserve β-cell function.

## RESULTS

### Palmitate inhibits autophagic flux and causes accumulation of autophagosome in pancreatic islets

To investigate the impact of saturated fatty acids on autophagic activity in pancreatic β-cells, we conducted experiments using isolated mouse islets or MIN6 insulinoma cells treated with palmitate (350 μM). The treatment of islets with palmitate resulted in ER stress, evidenced by dilated ER cisternae, the accumulation of autophagic vesicles, and reduced number of insulin granules, as visualized using transmission electron microscopy (Figure 1A). This accumulation indicated a disruption of autophagy by lipotoxicity in islets. In order to confirm the defect in autophagy, we examined levels of the autophagic markers LC3 and p62, and the autophagic flux using MIN6 cells stably expressing LC3-RFP-GFP. The protein amount of LC3II and p62 were found to be elevated (Figure 1B), and the ratio of yellow to red puncta was increased, indicating defective autophagic flux (Figure 1C).

**Figure 1.**
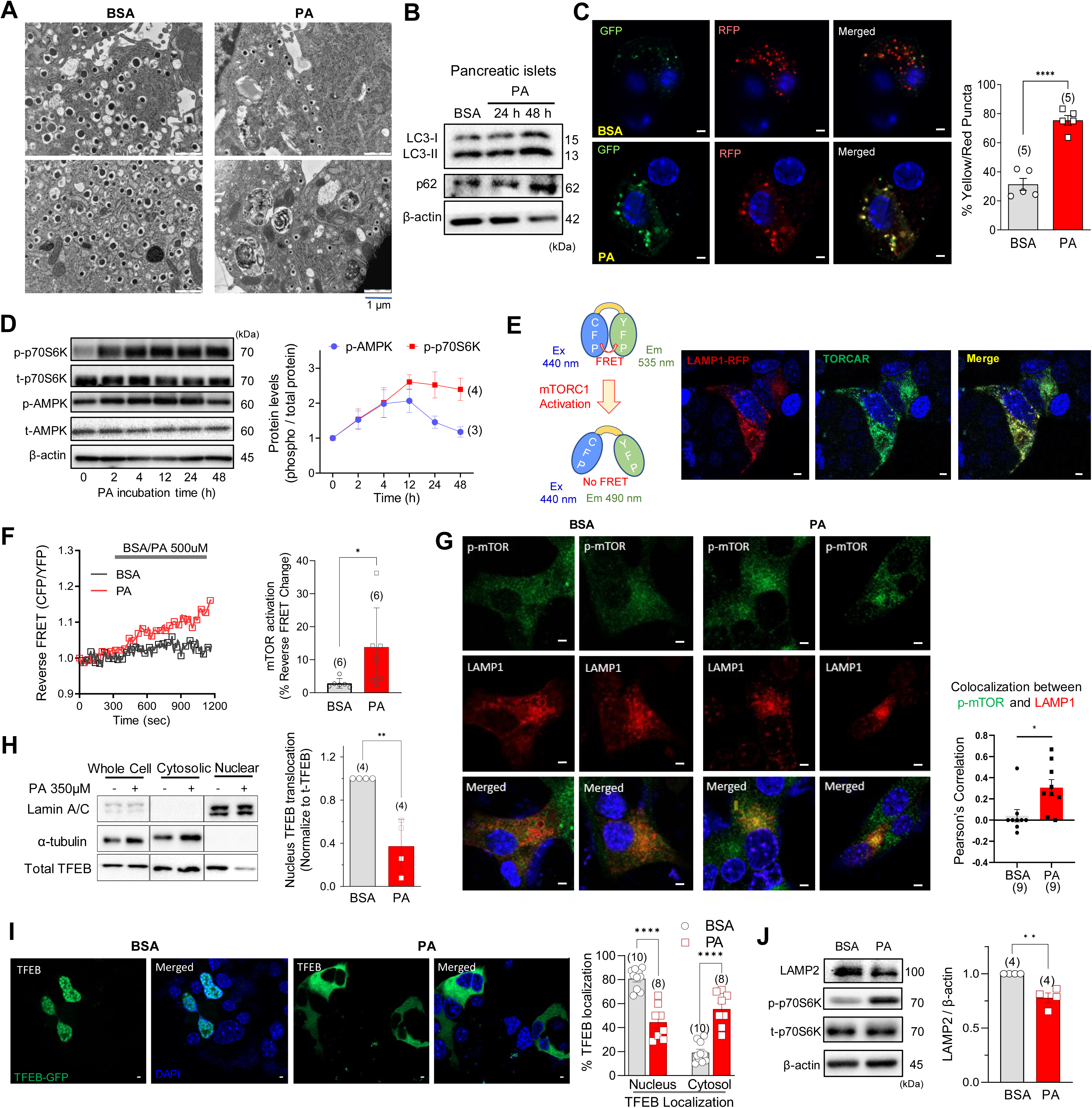
Palmitate induces autophagy defects mediated by mTORC1 activation at lysosomal membranes. (**A**) Representative pictures of electron microscopy showing that exposure to palmitate (PA; 350μM for 24h) causes accumulation of undigested autophagosomes, and dilated ER lumen, and reduces number of insulin granules in mouse pancreatic β-cells. (**B**) LC3-II and p62 accumulation by PA in pancreatic islets. (**C**) Elevated yellow puncta induced by PA in LC3-GFP-RFP-expressing MIN6 cells. (**D**) Time kinetics of p70S6K and AMPK phosphorylations by PA. (**E**) Lysosomal localization of TORCAR, an indicator of mTORC1 activity. (**F**) PA activates mTORC1 measured by TORCAR. (**G**) Colocalization of phosphorylated (activated) mTOR with LAMP1, a lysosome marker. (**H & I**) Decreased nuclear translocation and increased cytosolic localization of TFEB by PA. (**J**) Reduced LAMP2 by PA in MIN6 cells. Data are presented as means ± standard errors or deviations and (n) is the number of independent experiments except those in (H) which is the number of analyzed cells from at least 3 independent experiments. *p < 0.05; **p < 0.01; ****p < 0.0001.

Since both mTOR and AMP-activated protein kinase (AMPK) are known to be master regulators of autophagy, we assessed the activities of mTORC1 and AMPK during palmitate incubation. We observed that palmitate stimulated both AMPK and mTORC1 activity albeit with different kinetics; AMPK phosphorylation was transient returning to basal within 24 hours whereas mTORC1 activation was sustained even after 2 days of exposure to lipotoxic conditions (Figure 1D). The continued mTORC1 activation could be responsible for the observed inhibition of autophagy induced by palmitate(Nguyen *et al*., 2024). To monitor the kinetics of mTORC1 activation, we utilized the TORCAR biosensor, a fluorescence resonance energy transfer (FRET)-based reporter that detects phosphorylation of 4E-binding protein 1 (4EBP1)(Zhou *et al*, 2015). TORCAR fluorescence completely colocalized with lysosomal-associated membrane protein 1 (LAMP1), a lysosomal membrane protein (Figure 1E), and palmitate increased reverse FRET ratio of TORCAR, reflecting mTORC1 activity (Figure 1F).

Previous studies have indicated that the activation of mTORC1 occurs when it is recruited to the lysosomal surface in response to an abundance of amino acids(Sancak *et al*, 2010). Here, we observed that palmitate increased colocalization of activated (phosphorylated) mTORC1 with LAMP1, which suggests activation of mTORC1 on the lysosomal membrane (Figure 1G). Activation of mTORC1 negatively regulates autophagy through various mechanisms including the inhibition of lysosomal biogenesis by phosphorylating the transcription factor TFEB(Vega-Rubin-de-Celis *et al*, 2017). This phosphorylation leads to the retention of TFEB in the cytosol and hinders its translocation into the nucleus(Martina *et al*, 2012). To test whether palmitate inhibits the translocation of TFEB into the nucleus, MIN6 cells were treated with palmitate and examined for the localization of TFEB in cytosolic and nuclear fractions. As shown in Figure 1H, TFEB in the nuclear fraction was significantly reduced by palmitate incubation. Consistent with these findings, transiently expressed TFEB-GFP was retained in the cytosol of palmitate-treated MIN6 cells (Figure 1I). These findings indicate that palmitate inhibits TFEB activation. As a result, lysosomal biogenesis is impaired as also demonstrated by the reduced expression of the lysosomal marker LAMP2 (Figure 1K).

### Palmitate inhibits TRPML1-mediated lysosomal Ca^2+^ release

Transient Ca^2+^ increases in the perilysosomal microdomain play a crucial role in the fusion between autophagosomes and lysosomes, a critical step for the degradation of the autophagosome(Xu & Ren, 2015). Saturated fatty acids, such as palmitate, induce oxidative stress and disrupt organellar Ca^2+^ homeostasis(Ly *et al*., 2017), which may impair the autophagic process due to perilysosomal Ca^2+^ disturbances. To investigate this hypothesis, we analyzed the transcriptional levels of TRPML channels, the primary mediators of Ca^2+^ release from the lysosomal lumen. TRPML1 was the predominant channel expressed in both mouse pancreatic islets (Figure 2A) and MIN6 cells (Figure 2B). In order to examine the clinical relevance of TRPML1 expression in human islets, we conducted a meta-analysis comparing TRPML1 transcript levels in pancreatic islets from control and type 2 diabetes patients. As shown in the forest plot (left) and heatmap (right), TRPML1 expression was significantly lower in diabetic islets, similar to the pattern observed for TRPML3, but not for TRPML2 (Figure 2C).

**Figure 2.**
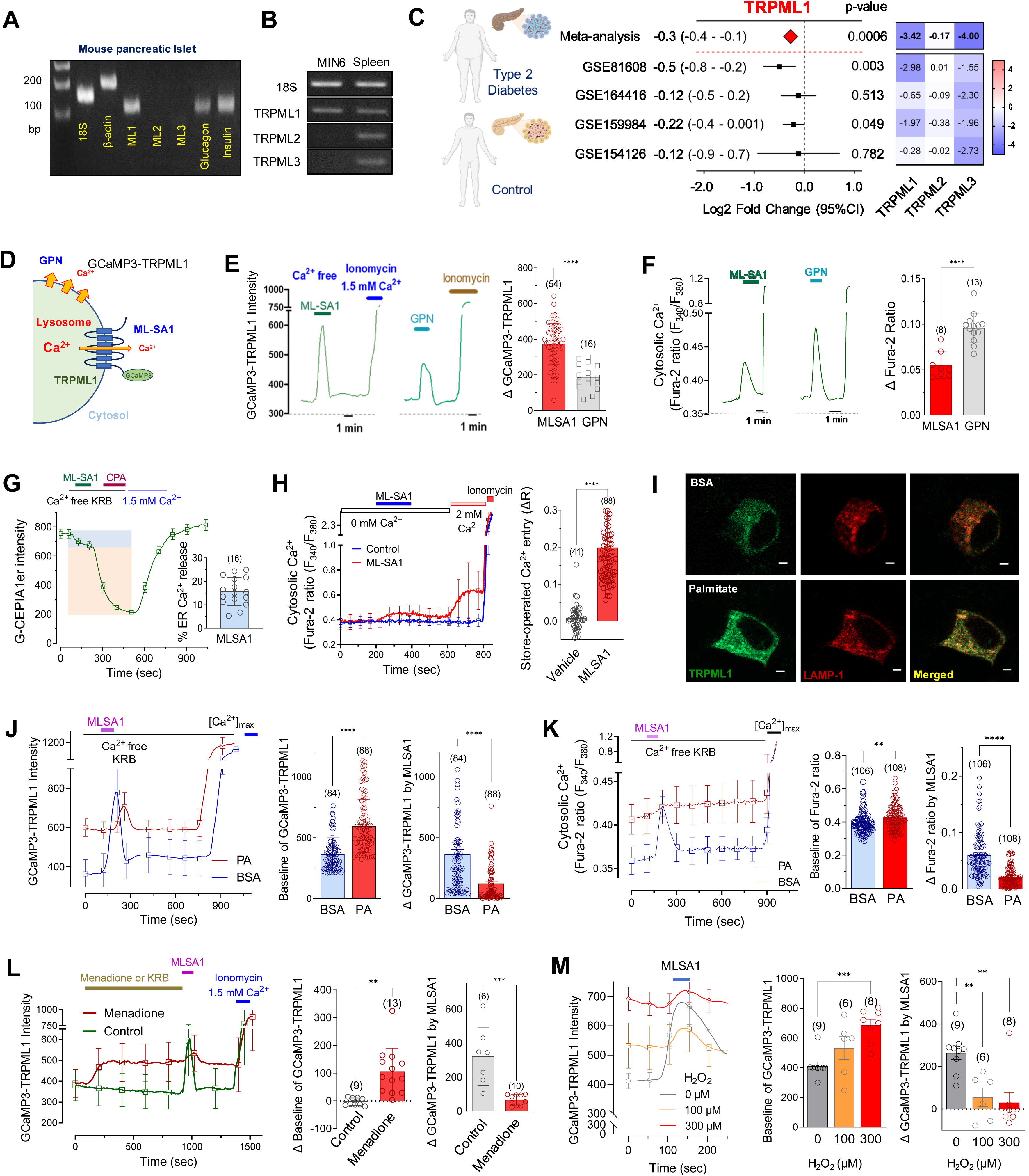
Palmitate elevates peri-lysosomal Ca^2+^ level and inhibits TRPML1-mediated lysosomal Ca^2+^ release. (**A & B**) Expression profile of TRPML channels in mouse pancreatic islets (A) and MIN6 cells (B). (**C**) Lower expression of TRPML1 in pancreatic islets from type 2 diabetes patients compared to control subjects in separate public datasets (black square) and their meta-analyzed fold change between control and diabetes patient (red diamond) calculated by IVW method. Heatmaps of TRPMLs show the z-values (top) calculated by meta-analysis and t-values from each dataset by DESeq2 analysis. (**D**) GCaMP3-TRPML1 fluorescent probe detecting perilysosomal Ca^2+^ level. (**E & F**) MLSA1 (10μM), a TRPML agonist, and GPN (100μM), a lysosomotrope, elevated perilysosomal (GCaMP3-TRPML1) and cytosolic (Fura-2) Ca^2+^ levels in MIN6 cells. (**G**) MLSA1 decreased ER Ca^2+^ content measured by G-CEPIA1er, suggesting ER Ca^2+^ release. (**H**) Cytosolic Ca^2+^ rose upon extracellular Ca^2+^ supplementation in MLSA1-treated group, indicating store-operated Ca^2+^ entry. (**I**) Fluorescence imaging of MIN6 cells expressing GCaMP3-TRPML1 treated with BSA or palmitate (PA) (**J & K**) PA elevated the basal level but abolished MLSA1 response of perilysosomal (J) or cytosolic (K) Ca^2+^ in MIN6 cells. (**L & M**) The oxidants, menadione (100μM) (L) or H_2_O_2_ (M), like PA, elevated the baseline Ca^2+^ and abolished the MLSA1 response. [Ca^2+^]_max_ implies cells in normal Ca^2+^ KRB with ionomycin. Data are presented as means ± standard deviations and (n) is the number of analyzed cells from more than 3 independent experiments, except those in L and M (from 2-3 independent experiments). **p < 0.01; ***p < 0.001; ****p < 0.0001.

To study Ca^2+^ signaling via TRPML1, three molecular probes were employed: Fura-2 to investigate cytosolic Ca^2+^, G-CEPIA1er to analyze ER Ca^2+^, and GCaMP3-TRPML1 fusion protein to measure perilysosomal Ca^2+^ levels (Figure 2D). Two specific drugs, MLSA1 (a TRPML1 agonist) and Gly-Phe β-naphthylamide (GPN, a lysosomotropic agent), were utilized to examine lysosomal Ca^2+^ signaling. In cells transfected with GCaMP3-TRPML1, MLSA1 (10μM) elicited a robust Ca^2+^ signal compared to a lesser increase induced by GPN (100μM) (Figure 2E). Using Fura-2 to measure changes in cytosolic Ca^2+^, it was observed that MLSA1 induced a smaller increase in total cellular Ca^2+^ level compared to the GPN-induced rise (Figure 2F). Furthermore, previous studies have suggested a close interaction between the ER and lysosomal Ca^2+^, which may be required for replenishing lysosomal Ca^2+^ content from the ER(Kang *et al*, 2024; Park & Lee, 2022). We found that MLSA1-mediated lysosomal Ca^2+^ release decreased the ER Ca^2+^ pool (∼15%), suggesting ’lysosomal Ca^2+^-induced ER Ca^2+^ release’ (Figure 2G). In response to depletion of the ER Ca^2+^ pool, store-operated Ca^2+^ entry (SOCE) was observed in MLSA1-treated cells. This substantiates the link between ER Ca^2+^ release and TRPML1-mediated lysosomal Ca^2+^ transient (Figure 2H). To explore whether this perilysosomal Ca^2+^ signaling occurs in other cell types, we measured fluorescence changes in human embryonic kidney (HEK) cells stably expressing GCaMP3-TRPML1. Similar to MIN6 cells, MLSA1 induced Ca^2+^ transients in HEK cells, followed by SOCE only in the MLSA1-treated group (Figure EV1).

To test our hypothesis regarding the effect of palmitate on lysosomal Ca^2+^ signaling, MIN6 cells expressing GCaMP3-TRPML1 were incubated with palmitate or bovine serum albumin (BSA) for 24 hours. Using fluorescence microscopy, we observed that palmitate-treated cells exhibited a significant increase in the GCaMP3 fluorescence signal, indicating elevated Ca^2+^ levels in the perilysosomal regions (Figure 2I). Importantly, MIN6 cells incubated with palmitate showed a diminished Ca^2+^ response to MLSA1, suggesting inhibition of Ca^2+^ release via TRPML1 channels (Figure 2J). Furthermore, palmitate-exposed cells displayed an elevated basal cytosolic Ca^2+^ and an attenuated MLSA1-induced Ca^2+^ response (Figure 2K). It has been reported that ROS triggers TRPML1 channel opening(Zhang *et al*., 2016). To compare the impact of palmitate on lysosomal Ca^2+^ regulation with that of ROS, MIN6 cells expressing GCaMP3-TRPML1 were exposed to the oxidizing agents, menadione (100μM) or H_2_O_2_ (100 and 300μM), and perilysosomal Ca^2+^ levels were recorded. Both menadione (Figure 2L) and H_2_O_2_ (Figure 2M) prominently elevated the basal fluorescence intensity, indicating an increase in perilysosomal Ca^2+^ levels. As a consequence, the Ca^2+^ response upon MLSA1 treatment was abrogated. These alterations in lysosomal Ca^2+^ signaling induced by ROS were similar to those caused by palmitate, suggesting that palmitate-induced oxidative stress may contribute to the pathologic dysregulation of lysosome and autophagic defects.

### mTORC1 activation prevents TRPML1-mediated lysosomal Ca^2+^ release and autophagic flux

As shown in Figure 1, palmitate activated mTORC1 located at the lysosomal membrane, interfering with nuclear TFEB translocation and lysosomal biogenesis. To understand the molecular mechanisms of mTORC1-mediated autophagy defects, we utilized MHY1485, an mTORC1 activator. As expected, MHY1485 (10μM) induced phosphorylation of S6K, accumulation of the autophagic marker p62, and increased cleaved caspase 3 in a time-dependent manner (Figure 3A). Additionally, the mTORC1 activity reporter TORCAR detected a marked increase in mTOR activity upon MHY1485 treatment (Figure 3B). Intriguingly, MHY1485 incubation for 12 h almost completely suppressed the MLSA1-induced perilysosomal Ca^2+^ response (Figure 3C), consistent with palmitate-suppressed TRPML1 activity (Figure 2J). Furthermore, when mouse islets were cultured with MHY1485, we observed enlarged ER lumen and the accumulation of autophagic vesicles (Figure 3D), similar to the phenotype of palmitate-treated islets (Figure 1A). Excessive mTORC1 activation by MHY1485 also led to mitochondrial membrane depolarization in MIN6 cells (Figure 3E). Co-incubation of palmitate with MHY1485 resulted in aggravated lipotoxicity in MIN6 cells, highlighting the detrimental consequences of mTORC1 activation in lipotoxicity (Figure 3F).

**Figure 3.**
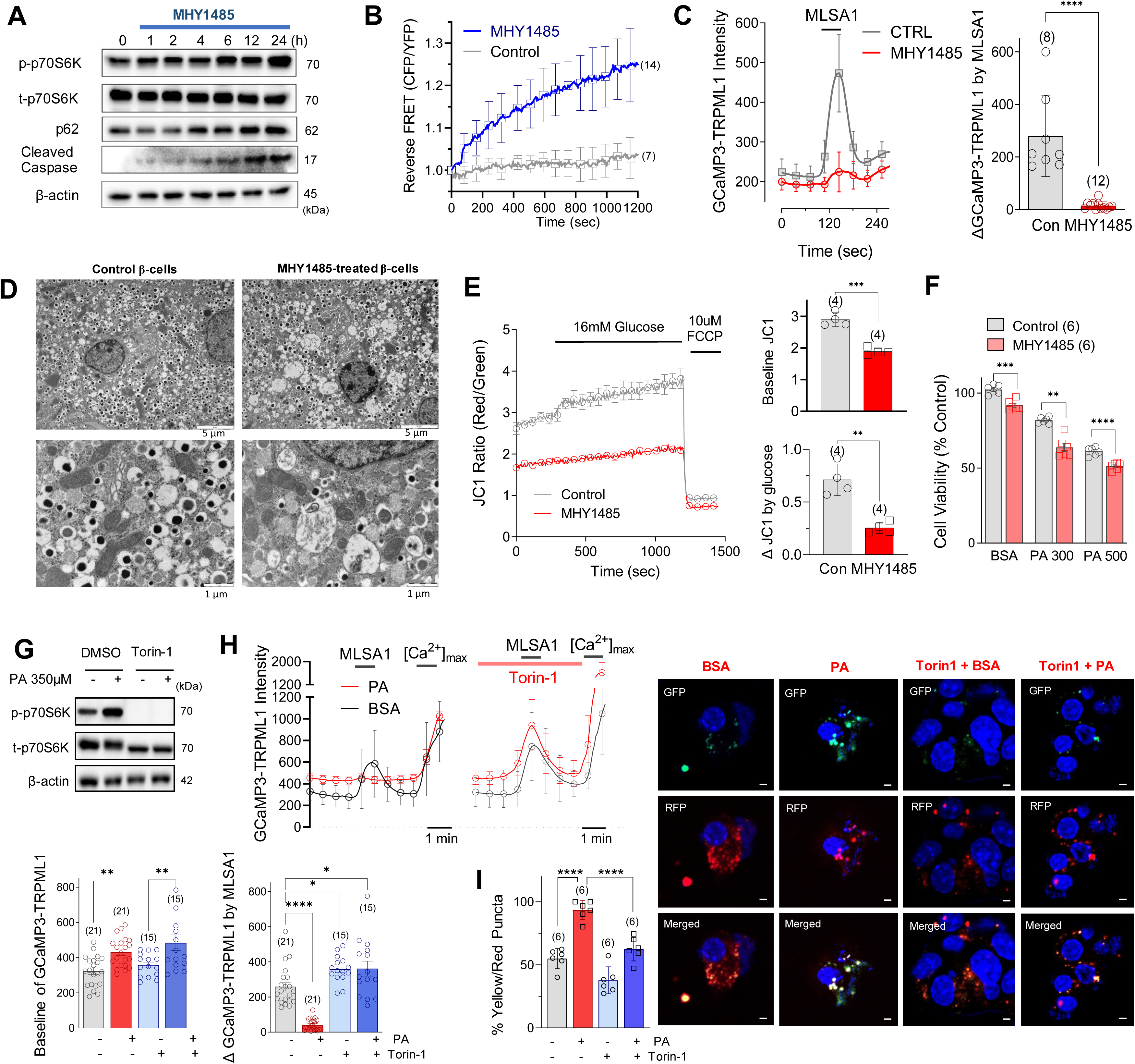
Inhibiting mTORC1 abolishes palmitate-induced TRPML1 suppression and autophagy defects. (**A & B**) mTORC1 activator, MHY1485 (10 μM), augmented mTORC1 signaling and increased p62 and cleaved caspase. (**C**) Pharmacologic activation using MYH1485 inhibited MLSA1-triggered lysosomal Ca^2+^ release in MIN6 cells. (**D & E**) mTOR activator caused accumulation of undigested autophagosomes and depolarized mitochondrial membrane potential leading to impaired glucose responses measured by JC1 fluorescence. (**F**) The mTOR activator, MYH1485, aggravated PA-induced cell death measured by using MTT assay (**G & H**) Torin-1 (100nM), an mTORC1 inhibitor, reduced basal perilysosomal Ca^2+^ level but recovered MLSA1-triggered lysosomal Ca^2+^ release. (**I**) Torin-1 recovered PA-increased yellow/red puncta in LC3-GFP-RFP expressing cells. Data are presented as means ± standard deviations (B,C,H) or errors (E,F) and (n) is the number of analyzed wells or cells from more than 3 independent experiments. *p < 0.05; **p < 0.01; ***p < 0.001; ****p < 0.0001.

To investigate the putative therapeutic value, we employed the specific mTORC1 inhibitor, Torin-1 (100nM), which completely abolished S6K phosphorylation (Figure 3G). Torin-1 restored perilysosomal Ca^2+^ signaling blunted by palmitate (Figure 3H). Additionally, using LC3-RFP-GFP, we observed that the inhibition of mTORC1 recovered autophagic flux in palmitate-treated cells (Figure 3I). Collectively, these findings provide evidence for the negative influence of mTORC1 activation induced by palmitate incubation on TRPML1 activity and its implications in autophagy and lipotoxicity.

### Scavenging mitochondrial ROS relieves palmitate-induced autophagic defects

In the previous section, we presented the association between oxidative stress and elevated perilysosomal Ca^2+^ levels, as well as the impaired response to MLSA1 (Figure 2L and 2M). To examine the potential impact of increased ROS on mTORC1 activation, we applied oxidative stress by exposing MIN6 cells to menadione (100μM) and assessed mTORC1 activation. Menadione strongly activated mTORC1 as demonstrated by phosphorylation of S6K (Figure 4A). Menadione also increased the reverse FRET ratio of TORCAR (Figure 4B). To study the role of mitochondrial ROS in palmitate-induced oxidative stress, we incubated dispersed mouse islet cells with mitoTEMPO (100nM), a mitochondrial superoxide scavenger. The application of mitoTEMPO significantly reduced both cytosolic and mitochondrial ROS levels, as measured by DCF and mitoSox, respectively (Figure 4C and 4D). Western blot analysis further revealed that mitoTEMPO blunted palmitate-induced S6K phosphorylation, LC3-II and p62 accumulation, as well as caspase activation (Figure 4E). Importantly, scavenging mitochondrial superoxide with mitoTEMPO prevented palmitate-induced perilysosomal Ca^2+^ dysregulation. mitoTEMPO lowered basal Ca^2+^ levels and recovered MLSA1-induced perilysosomal Ca^2+^ rises (Figure 4F). Furthermore, mitoTEMPO promoted TFEB translocation to the nucleus rescuing TFEB from palmitate-induced retention in the cytosol (Figure 4G). As a consequence of the normalization of lysosomal Ca^2+^ signaling, autophagic flux was restored, as measured using LC3-GFP-RFP processing (Figure 4H). These findings provide evidence that mitochondrial oxidative stress caused by palmitate induces mTORC1-mediated peri-lysosomal Ca^2+^ dysregulation and autophagic defects.

**Figure 4.**
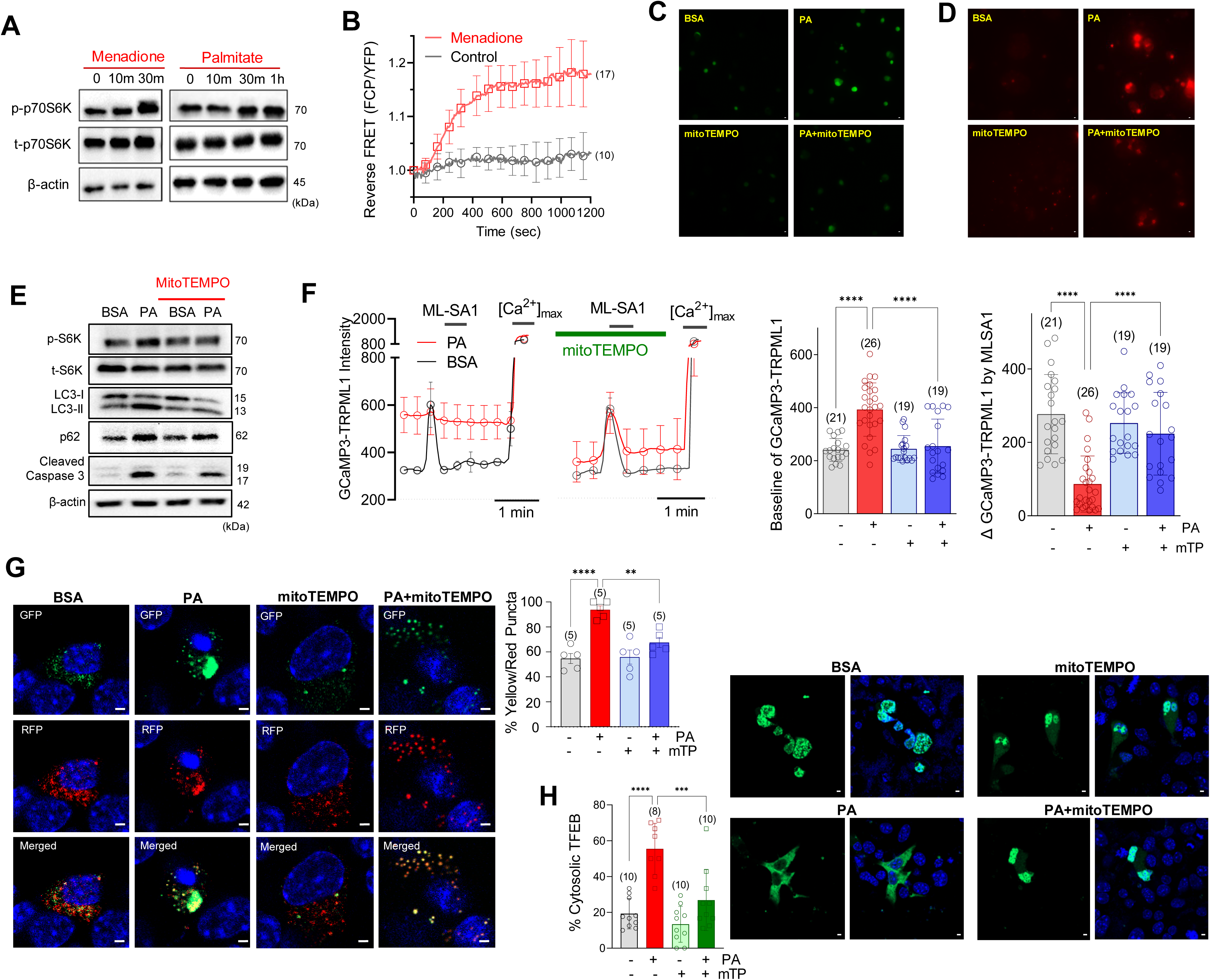
Scavenging mitochondrial superoxide prevents palmitate-induced TRPML1 inhibition and autophagy defects. (**A & B**) Menadione, an oxidant, activated mTORC1 signaling. (**C & D**) Reduction of cytosolic (DCF) and mitochondrial (mitoSox) ROS by the mitochondrial superoxide scavenger, mitoTEMPO (mTP, 100nM). (**E**) MitoTEMPO alleviated PA-increased LC3-II, p62, activated mTOR, and cleaved caspase. (**F**) MitoTEMPO prevented perilysosomal Ca^2+^ elevation and restored the MLSA1 response attenuated by PA. (**G**) MitoTEMPO counteracted PA-induced cytosolic retention of TFEB. (**H**) MitoTEMPO restored PA-induced defective autophagic degradation measured in LC3-GFP-RFP expressing MIN6 cells. Data are presented as means ± standard errors (G,H) or deviations (B,F). (n) is the number of independent experiments (G,H) or analyzed cells (B,F) from more than 3 independent experiments. **p < 0.01; ***p < 0.001; ****p < 0.0001.

### Preventing Ca^2+^ overload alleviates palmitate-induced mTORC1 activation and autophagic blockage

We have shown above that oxidative stress elicits strong mTORC1 activation (Figure 3A and 3B), which leads to detrimental consequences on lysosomal Ca^2+^ signaling and autophagic flux. However, the molecular connection between oxidative stress and mTORC1 activation remains unclear. Notably, it has been reported that amino acid-induced mTORC1 activation at the lysosomal membrane is dependent on Ca^2+^/Calmodulin (CaM)(Li *et al*, 2016a). To understand the pathogenic role of cellular Ca^2+^ dysregulation by palmitate, MIN6 cells were subjected to different conditions, including nominally Ca^2+^ free medium and intracellular Ca^2+^ chelation with BAPTA-AM (10μM). To further investigate the downstream signaling, we also used W7 (10μM) and KN62 (20μM) to inhibit calmodulin (CaM) and CaM kinase II (CaMKII), respectively. All conditions with lowered cytosolic Ca^2+^ level or CaM/CaMKII inhibition abolished palmitate-induced mTORC1 activation, indicating the key role of Ca^2+^ and CaM in the activation of mTORC1 during lipotoxicity (Figure 5A∼5C).

**Figure 5.**
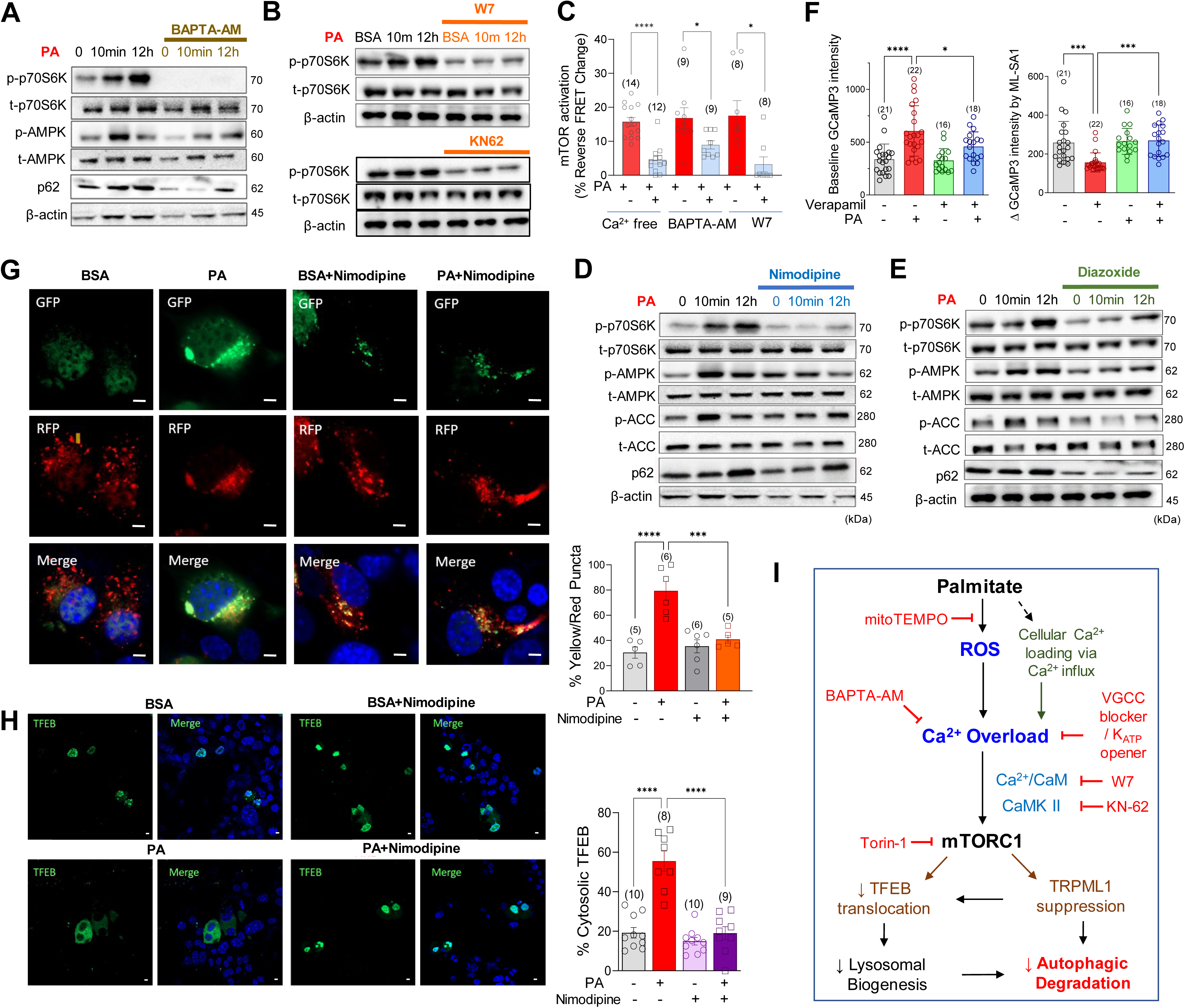
Reducing Ca^2+^ influx attenuates palmitate-induced mTORC1 activation and autophagy defects. (**A**) Intracellular Ca^2+^ chelation using BAPTA-AM inhibited PA-induced p70S6K and AMPK activation. (**B**) Inhibitors of calmodulin (W7, 10μM) and CaMKII (KN-62, 20μM) prevented PA-activated p70S6K. (**C**) PA-induced mTORC1 activation was abrogated by extracellular Ca^2+^ free medium, BAPTA-AM (10μM), or W7 measured in TORCAR-expressing MIN6 cells. (**D & E**) PA-induced activation of p70S6K and AMPK was abolished by nimodipine (D, 10μM), a VGCC inhibitor, or diazoxide (E, 200μM), an ATP-sensitive K^+^ channel opener. (**F**) PA effects on perilysosomal Ca^2+^ level and MLSA1 responses were prevented by a voltage-gated Ca^2+^ channel (VGCC) inhibitor, verapamil (10μM). (**G**) Nimodipine restored nuclear localization of TFEB. (**H**) PA-induced defective autophagic degradation was reestablished by nimodipine, measured in LC3-GFP-RFP expressing cells. (**I**) Hypothetical mechanisms of autophagic defects in PA-induced β-cell lipotoxicity. Data are presented as means ± standard errors (G,H) or deviations (C,F). (n) is the number of independent experiments (G,H) or analyzed cells (C,F) from more than 3 independent experiments. *p < 0.05; ***p < 0.001; ****p < 0.0001.

During β-cell metabolic activation, ATP-sensitive K^+^ (K_ATP_) channels close causing plasma membrane depolarization. As a consequence, voltage-gated Ca^2+^ channels (VGCCs) open and act as the primary Ca^2+^ influx mechanism triggering insulin secretion. To evaluate the impact of extracellular Ca^2+^ influx on palmitate-induced mTORC1 activation and autophagy suppression, MIN6 cells were incubated with either the VGCC blockers, verapamil (10μM) and nimodipine (10μM), or the K_ATP_ channel opener, diazoxide (200μM). Diazoxide induces strong hyperpolarization and prevents opening of VGCCs in insulin-secreting β-cells. Not only VGCC blockade but also K_ATP_ channel opening abrogated palmitate-induced mTORC1 activation and p62 accumulation, further confirming the critical role of VGCCs in palmitate-induced Ca^2+^ overload (Figure 5D and 5E). Verapamil recovered peri-lysosomal Ca^2+^ homeostasis, reduced basal Ca^2+^ level, and restored the Ca^2+^ response to MLSA1 (Figure 5F). This is consistent with our previous report about verapamil-induced suppression of oxidative stress and lipotoxicity(Ly *et al*, 2020). Moreover, inhibition of VGCC-mediated Ca^2+^ influx preserved autophagic flux under palmitate toxicity in MIN6 cells (Figure 5G), and restored the nuclear localization of TFEB (Figure 5H). Based on our findings, a molecular mechanism is proposed in which palmitate-induced cytosolic Ca^2+^ overload requires extracellular Ca^2+^ influx via VGCCs, which in turn activates mTORC1 through the involvement of CaM/CaMKII. mTORC1 activation suppresses TRPML1 activity and decreases TFEB translocation into the nucleus, ultimately contributing to the blockage of autophagy during lipotoxicity (Figure 5I).

### Activating the SERCA relieves palmitate-induced Ca^2+^ overload and rescues autophagic activity

ER Ca^2+^ depletion is a primary mechanism responsible for inducing ER stress and cytosolic Ca^2+^ overload in lipotoxicity. Multiple mechanisms contribute to the depletion of ER Ca^2+^ stores, and one of them is impaired smooth endo/sarcoplasmic Ca^2+^ ATPase (SERCA) activity(Ly *et al*., 2017). To probe for SERCA implication, we utilized CDN1163 (10μM), a SERCA pump activator, to investigate whether enhancing ER Ca^2+^ uptake through the SERCA pump could rescue autophagic impairment and cell death. To follow ER Ca^2+^ depletion, we monitored cytosolic Ca^2+^ increases triggered by SERCA pump inhibition using cyclopiazonic acid (CPA, 20μM). The incubation of cells with palmitate significantly diminished the content of ER Ca^2+^, while cotreatment with CDN1163 restored the ER Ca^2+^ pool depleted by palmitate (Figure EV2), which is consistent with our previous report(Nguyen *et al*, 2023). Furthermore, along with alleviating of ER stress, this SERCA activator reduced cytosolic and mitochondrial ROS generation caused by palmitate (Figure 6A and 6B). Palmitate-induced elevation of basal perilysosomal Ca^2+^ levels and inhibition of ML-SA1 responses were alleviated by co-treatment with CDN1163 (Figure 6C). Moreover, CDN1163 recovered palmitate-induced suppression of basal and maximal mitochondrial respiration, suggesting improvement of mitochondrial functions by activating ER Ca^2+^ uptake (Figure 6D). CDN1163 alleviated mTORC1 activation and p62 as well as LC3-II accumulation induced by palmitate (Figure 6E). Likewise, CDN1163 facilitated the translocation of TFEB into the nucleus and restored autophagic flux (Figure 6F and 6G). Consequently, CDN1163 inhibited cell death induced by palmitate in MIN6 cells (Figure 6H). By augmenting the activity of the SERCA pump, the uptake of Ca^2+^ into the ER lumen can restore ER Ca^2+^ levels and recover perilysosomal Ca^2+^ overload induced by palmitate. These actions could exert a beneficial effect on autophagic turnover by relieving cytosolic Ca^2+^ overload and ROS generation, which in turn mitigates mTORC1-mediated TRPML1 suppression in β-cells (Figure 6I).

**Figure 6.**
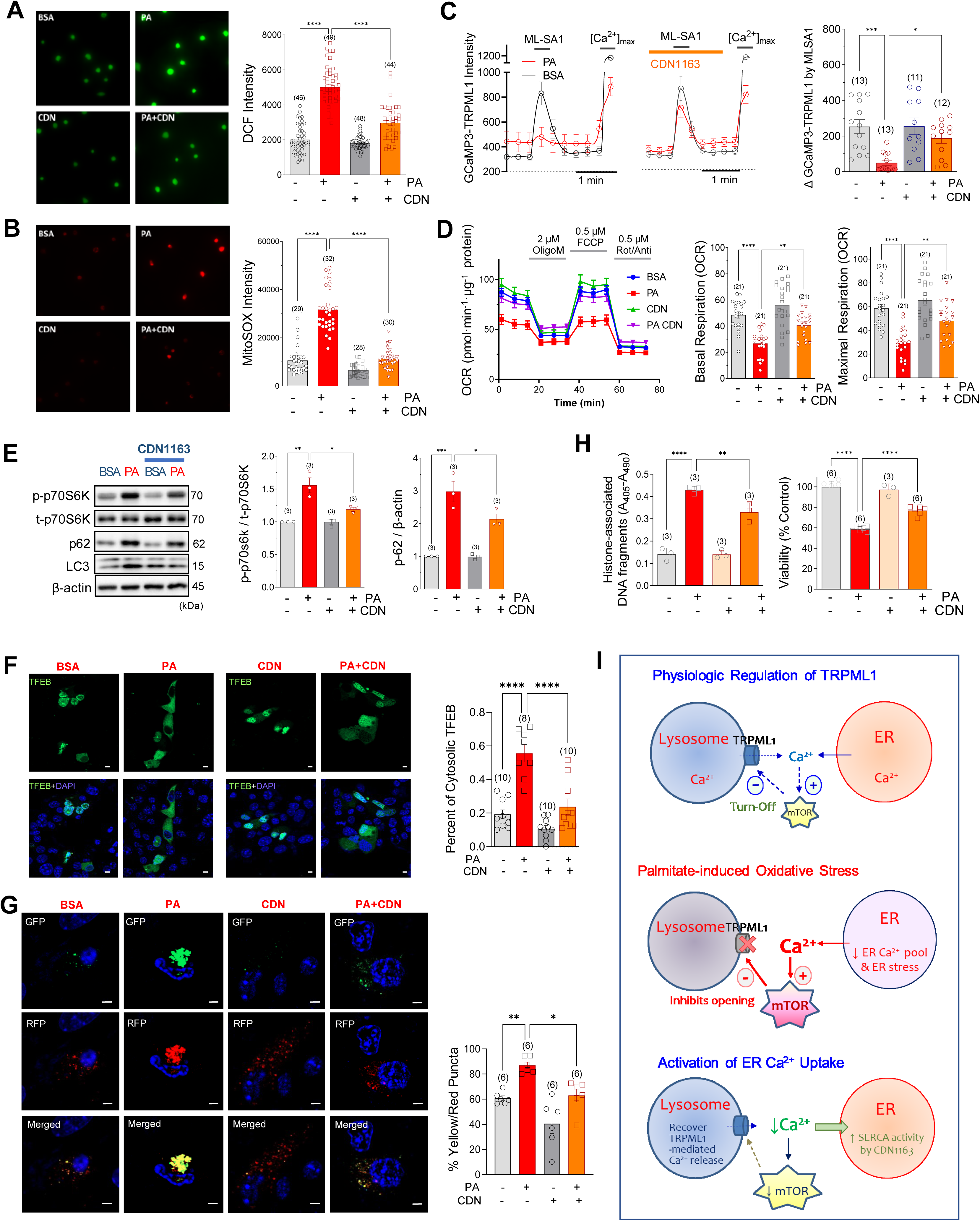
Activating ER Ca^2+^ uptake recovers peri-lysosomal Ca^2+^ homeostasis and autophagy activity. (**A & B**) CDN1163 (10μM) abrogated PA-induced cytosolic (DCF; A) and mitochondrial (mitoSOX; B) ROS generation. (**C**) CDN1163 prevented PA effects on perilysosomal Ca^2+^ level and MLSA1 responses. (**D**) CDN1163 recovered basal and FCCP-induced maximal oxygen consumption. (**E**) CDN1163 attenuated p70S6K activation and p62 accumulation. (**F**) CDN1163 prevented the inhibition of nuclear localization of TFEB.by PA (**G**) PA-induced defective autophagic degradation was recovered by CDN1163, measured in LC3-GFP-RFP expressing cells. (**H**) CDN1163 decreased PA cytotoxicity, measured by cell viability (MTT) and apoptotic (cell death detection) assays in MIN6 cells. (**J**) Schematic hypothesis of palmitate-induced TRPML1 suppression and its recovery by activating ER Ca^2+^ uptake.

## DISCUSSION

In the present study, we elucidated the mechanism underlying palmitate-evoked inhibition of autophagic flux in pancreatic β-cells. Importantly, palmitate induces prolonged activation of mTORC1 at the lysosomal membrane, which is driven by elevated peri-lysosomal Ca^2+^ levels. Dysregulated mTORC1 activation caused by palmitate disrupts physiologic Ca^2+^ conductance through lysosomal Ca^2+^ channels and prevents the nuclear translocation of TFEB. These disturbances result in significant pathogenic consequences that contribute to the blockage of autophagy. Our findings provide important insights into the critical role of oxidative stress-Ca^2+^ overload-mTORC1-linked cytopathology underlying the autophagic defects observed in pancreatic β-cells subjected to lipotoxicity.

Our previous investigations had focused on the depletion of ER Ca^2+^ and mitochondrial dysfunction associated with palmitate-induced oxidative stress(Ly *et al*., 2017; Xu *et al*, 2015). In order to attenuate mitochondrial superoxide production, we aimed to reduce the expression of mitochondrial Ca^2+^ uniporter (MCU), but observed palmitate-induced cytosolic Ca^2+^ overload and aggravation of autophagy blockage(Ly *et al*., 2020). These findings were corroborated by electron micrographic analyses, clearly showing extensive accumulation of undigested autophagosome-like vesicles in palmitate-incubated islets, which prompted the current study to understand the Ca^2+^ dynamics and homeostasis in inter-organellar ionic microdomains. We observed the retention of LC3-II and p62, as well as the downregulation of lysosomal biogenesis and function by palmitate underlining the importance of autophagosome degradation in β-cell lipotoxicity. To elucidate the molecular connection between palmitate and autophagic defects, we investigated the main regulatory signaling pathways in autophagy. Our results revealed that palmitate treatment activates the two antagonistic signaling events, AMPK and mTORC1, with different kinetics. AMPK activation was transient, whereas mTORC1 was sustained for an extended duration. The activity of mTORC1 is known to impede various phases of the autophagic process, through the regulation of key components such as ULK1(Jung *et al*, 2009), PIK3C3(Yuan *et al*, 2013), AMBRA1(Nazio *et al*, 2013), UVRAG(Kim *et al*, 2015), and TFEB(Martina *et al*., 2012). Thereby, activation of mTORC1 by palmitate ultimately leads to impaired autophagy and worsened cell fate. We also observed that palmitate markedly lowered the nuclear translocation of TFEB, possibly caused by mTORC1-mediated phosphorylation.

An intriguing observation from public datasets is that the transcript levels of the Ca^2+^ channels TRPML1 and TRPML3 are lower in pancreatic islets from T2D patients. Alterations in lysosomal Ca^2+^ channels expression and function in human diabetes pathogenesis has not been studied previously. In pancreatic islets, as well as MIN6 cells, the main isotype of TRPML channel was TRPML1 expressed in lysosomes. Of note, TRPML3 is less active at acidic pH and known to play a more prominent role in early endosomes(Kim *et al*, 2022). We investigated the role of the TRPML1 channel in organellar Ca^2+^ dynamics in β-cells and demonstrated that ER Ca^2+^ reservoir is utilized by the stimulation of TRPML1-mediated lysosomal Ca^2+^ release (Figure 2G). This was further supported by the activation of SOCE resulting from ER Ca^2+^ depletion (Figure 2H). Our findings are consistent with earlier reports that observed ER Ca^2+^ release (CICR) in response to Ca^2+^ release from lysosomes(Kilpatrick *et al*, 2016; Yuan *et al*, 2024). Two possible functional implications of CICR between lysosome and ER can be proposed. First, it has been suggested that the lysosomal Ca^2+^ pool may be replenished from the ER Ca^2+^ through physical contacts mediated by IP_3_ receptors. This ER to lysosome Ca^2+^ transfer is crucial for maintaining lysosomal Ca^2+^ homeostasis as well as for the autophagy process. Alternatively, the release of ER Ca^2+^ into the perilysosomal region may amplify the lysosomal Ca^2+^ signal, which is necessary for triggering the fusion between autophagosome and lysosome, thereby facilitating autophagic flux. To better understand these processes, further elucidation of the machineries involved in CICR is needed.

We have positioned the sustained activation of mTORC1 as a key mechanism linking lipotoxicity to autophagy defects. It has been proposed that TRPML1 and mTORC1 regulate each other through a feedback loop, which could enable transient regulation of TRPML1 during physiological stimulation(Sun *et al*, 2018). Our results show that prolonged pharmacological activation (Figure 3C) or chronic stimulation of mTORC1 during lipotoxicity (Figure 2J) negatively affects TRPML1 channel. Moreover, the detrimental consequences of pharmacological mTORC1 activation, including the accumulation of autophagosome, mitochondrial dysfunction, and cell death (Figure 3C & 3D), are qualitatively similar to those caused by palmitate (Figure 1). As anticipated, inhibition of mTORC1 using Torin-1 successfully restored TRPML1 activity and autophagic flux (Figure 3H & 3I). Based on this evidence we propose that mTORC1 could be a drug target to restore autophagic flux in β-cells suffering from lipotoxic stress.

We, then, elucidated the mechanisms linking lipotoxicity to mTORC1 activation. We observed that strong oxidants such as menadione and H_2_O_2_, elevated the basal peri-lysosomal Ca^2+^ levels, which may originate from both lysosomes and the ER. This elevation subsequently suppressed TRPML1 activity and the Ca^2+^ transient induced by MLSA1. Additionally, both oxidant agents and palmitate were found to induce sustained activation of mTORC1. Based on our findings, we postulate that palmitate-induced oxidative stress increases Ca^2+^ release into the peri-lysosomal microdomain where it is crucial for mTORC1 activation. This scenario was further validated by MitoTEMPO, a mitochondrial ROS scavenger, which effectively abrogated perilysosomal Ca^2+^ elevation and restored TRPML1 activity, TFEB nuclear translocation, and autophagic degradation in palmitate-treated cells. Oxidative stress is therefore likely causing peri-lysosomal Ca^2+^ overload, resulting in mTORC1 overstimulation. Previous studies have indicated that mTORC1 stimulation by nutrients, particularly amino acids, is mediated by Ca^2+^/CaM signaling, which is preceded by its recruitment to the lysosomal membrane(Li *et al*, 2016b; Sancak *et al*., 2010). Similar mechanisms may be responsible during palmitate-induced Ca^2+^ signaling, as demonstrated by our finding that palmitate activates mTORC1 on the lysosomal membrane (Figure 1G) that was abolished by intracellular Ca^2+^ chelation using BAPTA-AM (Figure 5A). Furthermore, we demonstrated that inhibitors of CaM (W7) or CaMKII (KN62) abolished mTORC1 activation induced by palmitate (Figure 5B), suggesting a Ca^2+^/CaM-dependent mTORC1 activation under lipotoxic conditions. It has been proposed that cytosolic Ca^2+^ overload is a crucial mechanism underlying β-cell glucolipotoxicity(Vogel *et al*, 2020). An uncontrolled increase in cytoplasmic Ca^2+^ has been known to trigger several deleterious effects, including activation of protein kinase C, c-Jun N-terminal kinase, CHOP, calpains, and caspases, ultimately leading to mitochondrial dysfunction, ER stress, and apoptosis(Carafoli, 2002; Cunha *et al*, 2008). Here, we propose another mechanism that sustained activation of mTOR and the resultant defective autophagy is an important part of the cytotoxic mechanisms associated with Ca^2+^ overload.

In order to reduce cellular Ca^2+^ content preventing Ca^2+^ overload, we applied different strategies, including the direct inhibition of VGCCs and the prevention of their opening through hyperpolarization, both of which decrease extracellular Ca^2+^ influx(Sargsyan *et al*, 2008). Consistent with the effects observed with BAPTA-AM treatment, the use of VGCCs’ blockers or K_ATP_ opener prevented mTORC1 activation and autophagy defects in the presence of palmitate. This suggests that counteracting cytosolic Ca^2+^ load holds promising therapeutic potential for mitigating β-cell lipotoxicity. Oxidative stress in the ER activates the release of Ca^2+^ from the ER and inhibits active Ca^2+^ reuptake via SERCA. We demonstrated that CDN1163, a SERCA activator, rescued pancreatic β-cells from lipotoxicity by enhancing ER Ca^2+^ uptake(Nguyen *et al*., 2023). In this investigation, we established that promoting ER Ca^2+^ uptake recovered intracellular Ca^2+^ homeostasis, improved mitochondrial function, as well as glucose-stimulated insulin secretion, and ultimately rescued cells from lipotoxicity-induced death. Additionally, CDN1163 restored autophagic flux in β-cells, along with the normalization of perilysosomal Ca^2+^ levels and TRPML1-mediated Ca^2+^ transient. These findings support the notion that promoting ER Ca^2+^ uptake can alleviate cytosolic Ca^2+^ overload, downregulate mTORC1 signaling, and restore autophagy in palmitate-induced lipotoxicity. Our data highlight the potential interplay between oxidative stress, Ca^2+^ dysregulation, and mTORC1 activation in the context of TRPML1 inhibition and defective autophagic degradation in β-cell lipotoxicity, as illustrated in Figures 5I and 6I.

Collectively, our study identifies a novel ROS-Ca^2+^ overload-mTORC1 cytopathic mechanism that underlies impaired autophagy and proposes various protective strategies to correct or relieve pathogenic alteration in β-cell lipotoxicity. Prolonged activation of mTORC1 by saturated fatty acids plays a critical role in inhibiting autophagy, as it suppresses TRPML1 activity and diminishes the nuclear localization of TFEB. Notably, reducing cellular Ca^2+^ loads by inhibiting extracellular Ca^2+^ influx or promoting Ca^2+^ reuptake into the ER could serve as efficient therapeutic strategies to alleviate oxidative stress and counteract uncontrolled mTOR activation. These results contribute to enhance our understanding of the mechanisms behind lipotoxicity-induced autophagic inhibition and may inform the development of therapeutic interventions aimed at restoring autophagy and preserving pancreatic β-cell function in lipotoxic states. Investigations into other pathological conditions characterized by the accumulation of undigested autophagosomes could broaden the applicability of the therapeutic strategies suggested by our results.

## METHODS

### Reagents and tools table

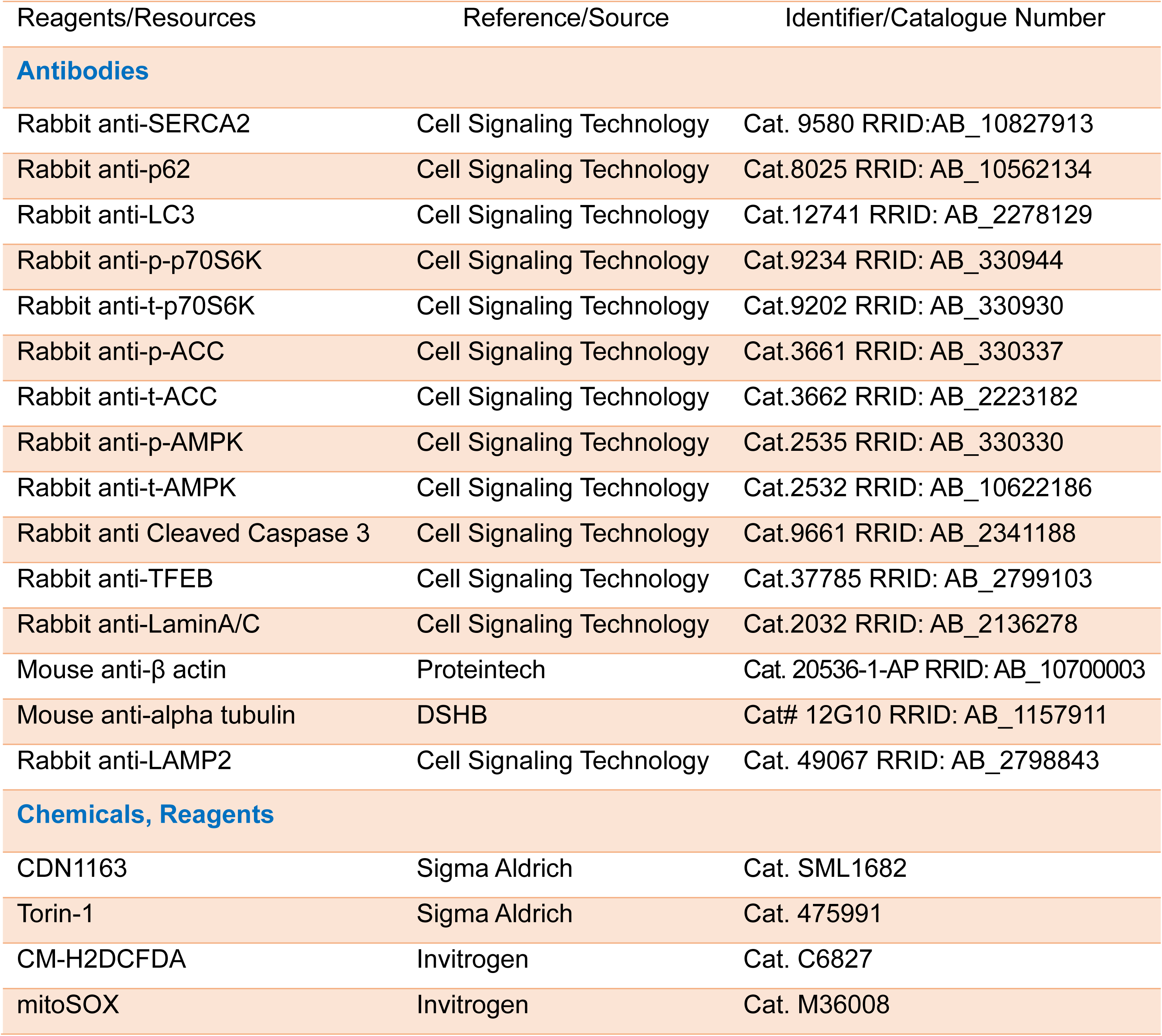
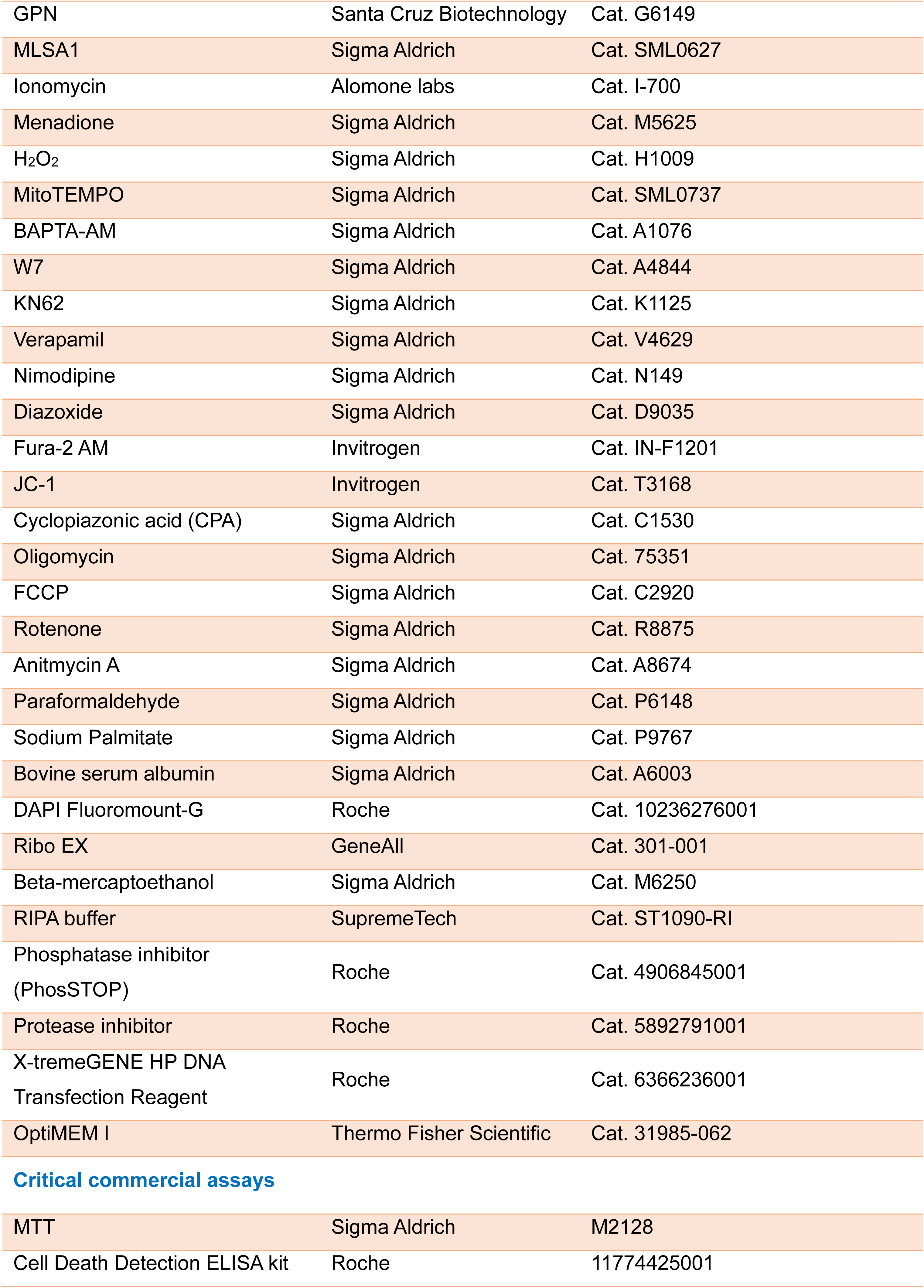
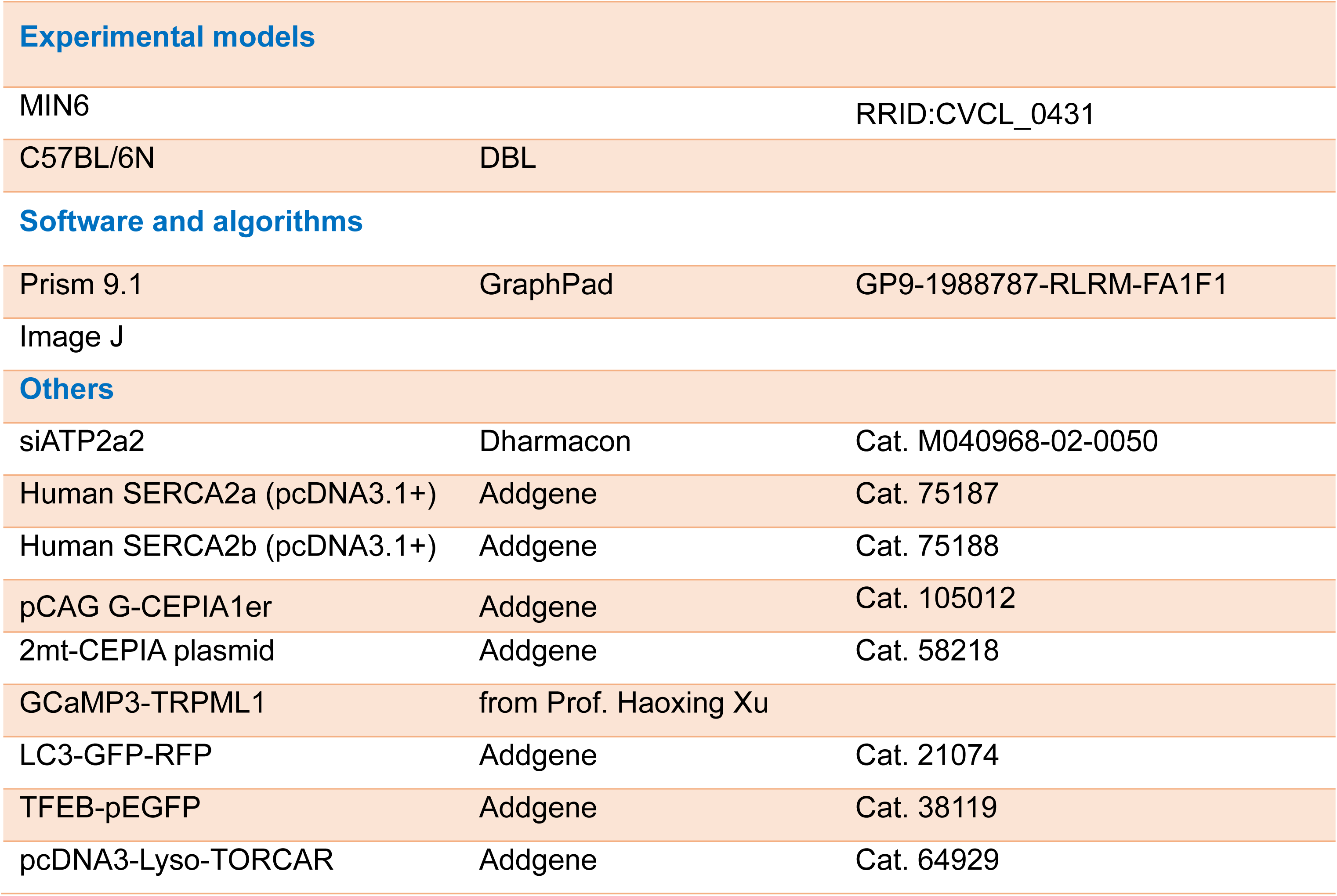

### Cell culture

MIN6 cells were cultivated in a controlled environment with 5% CO2 at 37°C, using Dulbecco’s modified Eagle’s medium supplemented with 12mmol/l glucose. The culture medium consisted of fetal bovine serum (10%), β-mercaptoethanol (5ul/l), streptomycin (100µg/ml), and penicillin (100U/ml) obtained from Hyclone, Thermo Fisher Scientific, Lafayette, CO. Passages ranging from 27 to 39 were selected for our investigation. The medium was refreshed every 2 days, and cells were passaged once a week following detachment using trypsin-EDTA. Prior to conducting experiments, the MIN6 cells were allowed to settle and cultured for 12-24 hours.

### Mouse islet isolation

All animal care and procedures were performed in accordance with the policies and approval (YWC-200907-2) of the Institutional Animal Care and Use Committee of Yonsei University Wonju College of Medicine. Ten to twelve-week-old C57BL/6N male mice (from DBL) were sacrificed for islet isolation.

A C57BL/6N male mouse was sacrificed by cervical dislocation after general anesthesia. After disinfection by 70% ethanol, the mouse was laid on the tray with tail down. A triangle was incised on the skin starting from the tail. Skin was rolled up and pulled up over the head of sternum, then hemostatic forceps were clamped at the sternum to keep the abdominal cavity opened. Guts and organs were displaced to the right side and the common bile duct was uncovered. By tracking along the common bile duct, ampulla of Vater could be found. It appeared as a white spot and should be clamped by hemostatic forceps. The two forceps were perpendicular. The common bile duct was then separated from vessels and liver and was catheterized using 5ml syringe with 30.5G needle. About 3ml of collagenase solution was injected into the common bile duct, distending the pancreas. Dressing forceps were used to detach guts from pancreas. The pancreas was cut off from spleen and was quickly put into 50ml conical tube with 5ml collagenase on ice.

The water bath (37°C) and cool centrifugation machine (4°C) should be turned on 1 h before the procedure. Tubes containing islets should be kept in water bath at 37°C for 20 min. Every 2-3 min, tubes were shaken vigorously and the pancreas gradually dissolves into fine particles. The incubation was terminated by adding 50ml of medium and centrifuging the tubes at 1500rpm at 4°C for 1min. Supernatant was removed, followed by 3 times washes. Then the pellet was suspended and transferred to a 15ml tube which was centrifuged at 1500rpm for 5 min at 4°C. Using Lympholyte to separate islets from the exocrine tissue by centrifuging at 1800 rpm for 15 min. Islets appeared as a ring between 2 different solutions. The islet layers were pipetted and transferred to two 50ml tubes which were centrifuged at 2000 rpm for 1min at 8°C. This wash was repeated 3 times. Thereafter, islets were resuspended and transferred to a petri dish and incubated at 37°C. Islets were maintained in RPMI 1640 supplemented with 10% FBS and 1% P/S.

### Cell viability (MTT) and apoptotic (cell death detection) assays

MIN6 cells were plated at 5×10^4^ cells/well in a 96 well plate. Drugs were pre-treated for 30 minutes before 24 hours co-incubation with 500uM palmitate or BSA. MTT solution was prepared by dissolving 3-[4,5-dymethylthiazol-2-yl]-2,5-diphenyl tetrazolium bromide in fresh DMEM medium at 0.5mg/ml. MTT assay was initiated by replacing culture medium with 100ul of MTT solution. The plate was put in an incubator during two hours. After careful suction of medium to avoid cell detachment, 100ul of DMSO was pipetted into each well to dissolve the formazan. The plate was then gently swirled for 10 minutes, and absorbance value of each well was measured at 570nm (reference wavelength 650nm) by Epoch^TM^ Microplate Spectrophotometer (Bio-Tek, Winooski, VT).

The amount of cytoplasmic mono- and oligonucleosome-associated histone-DNA complexes in cell lysates were quantified in a sandwich ELISA assay (Cell Death Detection ELISA Plus kit, Roche Diagnostics). 5×10^4^ MIN6 cells/well were seeded into 96 well plate. Drugs were added for 30 min before 24 hours co-incubation with 500uM palmitate or BSA. The assay was performed by adding 200ul of lysis buffer supplied in the kit to each well, and the plate was incubated for 30 minutes at room temperature. After centrifugation, 20ul of supernatant was collected, and the following steps were conducted following the protocol of the manufacturer. After 5-minute incubation with a peroxidase substrate, absorbance values were measured at 405nm (reference wavelength 490nm) in an Epoch^TM^ Microplate Spectrophotometer (Bio-Tek, Winooski, VT).

### Western blot analysis

For total protein extraction, cells seeded on 6-well plates were washed with ice-cold PBS and lysed with cold RIPA buffer (Thermo Fisher Scientific) containing protease inhibitor cocktail (Roche Diagnostics, Mannheim, Germany). The supernatants from lysates were loaded for SDS-PAGE and then transferred to polyvinylidene difluoride membrane (Merck Millipore, Billerica, MA). The membrane was blocked by 6% skimming milk for 1 hour at room temperature, followed by incubation with primary antibody at 4°C overnight. Horseradish peroxidase (HRP)-conjugated secondary antibody against either mouse or rabbit IgG was incubated for 1 hour at room temperature. The bands were visualized with a ChemiDocTM XRS+ imaging system (Bio-Rad) using enhanced chemiluminescence (ECL) solution (Luminata Forte, Millipore, Billerica, MA).

### Reverse transcription (RT) and real-time PCR

Total RNA was extracted from MIN6 or islets using Hybrid-R RNA kit. cDNA was synthesized from 0.5-1g RNA with qPCR RT Mastermix according to the manufacturer’s protocol. cDNA was amplified using Accupower HotStart PCR premix (Bioneer) to detect the expression of genes of interest. For real-time PCR, each cDNA sample was mixed with SYBR Green PCR mastermix in a triplicate manner in a real-time PCR machine. ΔΔCt method using *Rn18s* as an internal control was used to analyze relative expression.

### Cytosolic and mitochondrial ROS measurement

5×10^4^ cells were plated onto 12mm coverslips embedded in 4 well plate before treatment with Palmitate (350µM, 24 hours) or BSA. Cells were loaded with 2.5µM CM-H2DCFDA or 5µM mitoSOX for 10 min at 37°C to detect intracellular or mitochondrial ROS, respectively. Excitation/emission wavelengths for DCF and mitoSOX were 490/535 nm and 514/560nm, respectively. Fluorescence intensity was quantified using the IX81 inverted microscope with MetaMorph 6.1.

### mTORC1 activity measurement using TORCAR

MIN6 cells were plated onto 12mm coverslips. Next day, cells were transfected with TORCAR plasmid. After 48 hours MIN6 cells expressed TORCAR and were treated with or without inhibitors. Prior to the experiment, cells were serum-starved overnight and amino acid for 2 hours to reduce basal mTORC1 activity. Performing live-cell imaging using an IX-73 inverted microscope (Olympus, Tokyo, Japan). During the measurement, cells were stimulated by Palmitate (or BSA for vehicle), and the resulting changes in FRET were recorded. Reversed FRET was used to indicate the activation of mTORC1.

### Perilysosomal, ER, cytosolic Ca^2+^ measurement

3×10^4^ cells suspended in 500µl of complete medium were seeded onto 12mm coverslips embedded in 4 well plates before treatment with palmitate (350µM, 24h) or BSA. Cells were loaded with 1µM Fura-2 AM dissolved in KRBB for 30 min in an incubator. Cells were washed three times by KRBB. During recording of the Fura-2 signals, the chamber was perfused with Ca^2+^ free KRBB in the presence of 200µM sulfinpyrazone unless otherwise mentioned. The chamber was placed on the IX-73 inverted microscope platform (Olympus, Tokyo, Japan) attached to a complementary metal-oxide-semiconductor camera and a LED illuminator. The fluorophore was alternately excited at 340 and 380nm while emission was recorded at 510nm using MetaFluor 3.1. At the end of each experiment, 10µM ionomycin was added to induce a maximal ratio for comparison between control and palmitate treated cells.

To measure ER Ca^2+^ and lysosomal Ca^2+^, we used G-CEPIA1er (Addgene plasmid #105012) and GCaMP3-TRPML1 which is generously provided by Prof. Haoxing Xu, respectively, were introduced by transfection 48 hours prior to the experiment. The fluorescent emission at 535nm was captured and subsequently analyzed after excitation at 488nm.

### Immunostaining and confocal microscopy

MIN6 cells were subjected to transfection with LC3-GFP-RFP or TFEB-GFP constructs. Follo wing a two-day incubation period, the transfected cells were exposed to either palmitate or BSA, with or without additional drugs. Subsequently, the cells were fixed using 4% paraformaldehyde at room temperature in the absence of light for 15 minutes. Images were acquired utilizing a Zeiss LSM 800 confocal microscope and ZEN2.3 software. To quantify the yellow and red puncta in LC3-RFP-GFP, ImageJ software from NIH was employed. For the assessment of TFEB-GFP loc alization, the distribution of TFEB in the cytosol or nucleus was quantified.

### Public transcriptome dataset collection and Meta-analysis

A multidisciplinary team of physiologists, data scientists, and clinicians conducted a comprehensive literature and database review to identify candidate datasets, ultimately collecting full-transcriptome profiling datasets from human beta cells (GSE81608, GSE164416, GSE159984, GSE154126). These datasets were standardized through preprocessing, log transformation, and harmonization. Differential expression analysis between diabetes and control groups was performed using DESeq2, followed by a meta-analysis employing the Inverse-Variance Weighted (IVW) method implemented in the METAL software.

### Statistical analysis

All experiments were conducted with randomized samples and expressed as mean ± standard deviation (SD) or standard error of the mean (SEM). The data analyses were blinded and statistical analysis was undertaken only for studies where number of independent experiments per group was at least 5. Using Prism software (Version 9.1.0.221, GP9-1988787-RLRM-FA1F1, GraphPad Software, San Diego, CA, USA), unpaired two-tailed Student’s t-test, one-way or two-way analysis of variance (ANOVA) were performed. Post-hoc Tukey multiple comparison test after ANOVA were performed only if F achieves the statistical significance. A p value of less than 0.05 was considered statistically significant.

## Abbreviations

AMPK: AMP-activated protein kinase
BSA: bovine serum albumin
CaM: calmodulin
CaMKII: calmodulin-dependent protein kinase II
ER: endoplasmic reticulum
GPN: Gly-Phe β-naphthylamide
K_ATP_ channel: ATP-sensitive K^+^ (K_ATP_) channel
LAMP1: lysosomal-associated membrane protein 1
MCOLN: mucolipin TRP cation channel 1
mTORC1: Mechanistic target of rapamycin C1
PI3P: phosphatidylinositol 3-phosphate
ROS: reactive oxygen species
SERCA: smooth endo/sarcoplasmic Ca^2+^ ATPase
SOCE: store-operated Ca^2+^ entry
TFEB: nuclear transcription factor EB
TRPML1: transient receptor potential mucolipin 1
T2D: Type 2 diabetes
VGCCs: voltage-gated Ca^2+^ channels

## Author Contribution

Conceptualization: H.T.N., L.D.L., K-S.P.; Methodology: H.T.N., S.K.L., C.N-P, T.L., K-S.P.; Investigation, H.T.N., L.D.L., T.T.N., S.K.L., C.N-P., S.L.; Funding: K-S.P.; Original Draft Writing: H.T.N., K-S.P.; Review & Editing: H.T.N., T.L., S-K.C., M-S.L., A.W., C.B.W., K-S.P.

## Acknowledgements

This work was supported by the Medical Research Center Program (2017R1A5A2015369 and RS-2024-00409403) and Bio & Medical Technology Development Program (2020M3A9D8039920 and RS-2024-00397929) of the National Research Foundation of Korea (NRF) funded by the Korea government (MSIT).

---

**Figure EV1.**
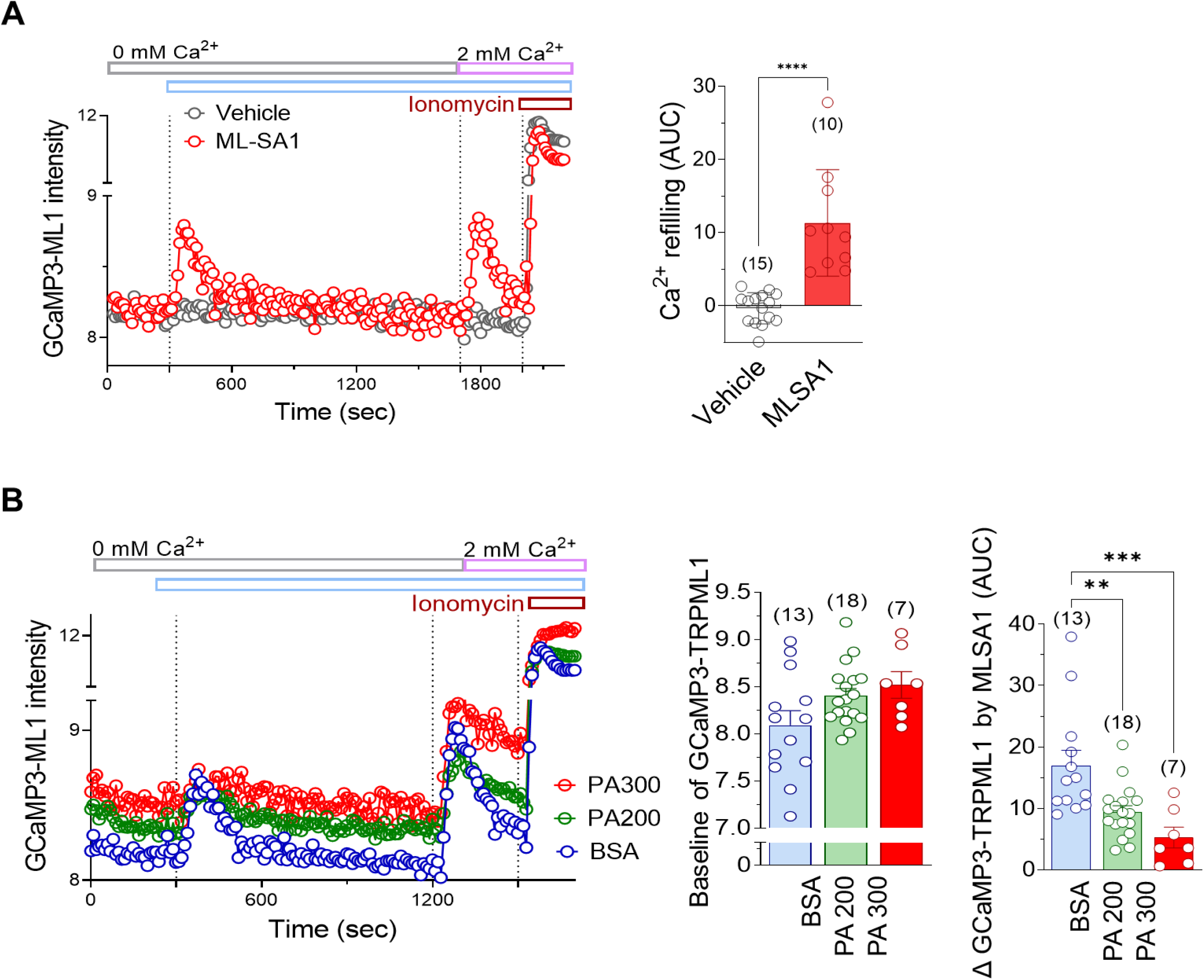
Perilysosomal Ca^2+^ measurement using high-throughput fluorescence reader with GCaMP3-TRPML1-expressing cells. (A) Perilysosomal Ca^2+^ rise upon extracellular Ca^2+^ addition, mediated by MLSA1-triggered SOCE. (B) Elevated the baseline level of perilysosomal Ca^2+^, but abolished MLSA1 response by palmitate (PA) in multi-well plates seeded with GCaMP3-TRPML1-expressing HEK cells. Fluorescence intensity from each well were measured by using FlexStation. (n) is the number of analyzed wells from at least 3 independent experiments. **p<0.01; ***p<0.001; ****p<0.0001.

**Figure EV2.**
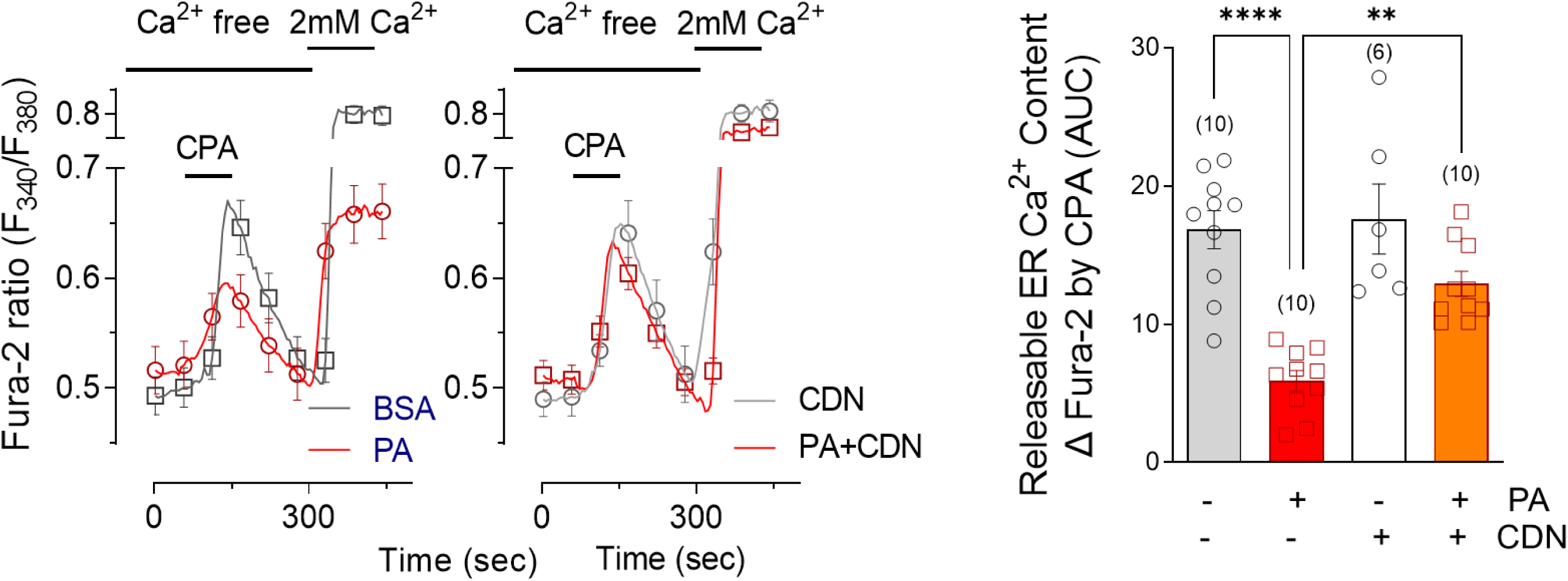
SERCA activator recovers ER Ca^2+^ depletion by palmitate. ER Ca^2+^ release due to cyclopiazonic acid (CPA) was reduced in palmitate-treated MIN6 cells, which was normalized by pretreatment with CDN1163, a SERCA activator. **p<0.01; ****p<0.0001.

